# On the efficacy of restoration in stream networks: comments, critiques, and prospective recommendations

**DOI:** 10.1101/611939

**Authors:** David Murray-Stoker

## Abstract

Swan and Brown (2017) recently addressed the effects of restoration on stream communities under the meta-community framework. Using a combination of headwater and mainstem streams, Swan and Brown (2017) evaluated how position within a stream network affected the outcome of restoration on invertebrate communities. Ostensibly, their hypotheses were partially supported as restoration had stronger effects in headwater streams: invertebrate taxonomic richness was increased and temporal variability decreased in restored reaches; however, these results were not consistent upon closer scrutiny for both the original paper (Swan and Brown 2017) and the later erratum (Swan and Brown 2018). Here, I provide a secondary analysis of the data, with hypotheses and interpretations in the context of stream, meta-community, and restoration ecology. Swan and Brown (2017, 2018) evaluated the effect of restoration on sites receiving various combinations of in-channel manipulation and riparian reforestation treatments. Given the difference in the relative importance of environmental filtering and dispersal between headwaters and mainstems and the structure of river networks, I contend that different restoration treatments have differential effects between headwaters and mainstems. I hypothesized in-channel manipulations would have more consistent effects between headwaters and mainstems compared to riparian reforestation, and I used this hypothesis to guide site selection in the re-analysis. I then compared results from the re-analysis to those presented by Swan and Brown (2017, 2018). I did not find any effects of restoration on local diversity, spatial dissimilarity, or temporal variability, let alone differential effects of restoration between headwaters and mainstems; these results are contrary Swan and Brown (2017, 2018), who reported that restoration increased taxonomic richness, increased spatial dissimilarity, and decreased temporal variability in restored headwater streams. I demonstrate further that the statistical tests conducted by Swan and Brown (2017, 2018) were invalid and, therefore, recommend the use of the results presented here. More broadly, I suggest, in agreement with Swan and Brown (2017, 2018) and a growing body of research, that river and stream restoration will likely have greater success if a regional approach is taken to designing and implementing restoration projects.

## Introduction

In a recent study, Swan and Brown (2017) evaluated how restoration affected community diversity in streams through the use of metacommunity theory. Under this framework, local effects are associated with species’ niches while regional effects are more associated with dispersal (Leibold et al. 2004). In the context of stream networks, headwaters are isolated patches more likely to be impacted by niches and environmental characteristics and mainstems are well-connected more likely to be affected by dispersal (Heino et al. 2003, Leibold et al. 2004, Grant et al. 2007, Altermatt 2013, Heino 2013). Restoration of stream habitats was therefore expected to have a greater impact on communities in headwaters relative to mainstems (Swan and Brown 2017).

Although Swan and Brown (2017, 2018) noted that restoration techniques can vary in intrusiveness on stream ecosystems, they did not account for this in their experimental design and statistical analyses. Restored streams in their study received various combinations of bank stabilization, in-channel manipulation, and riparian reforestation (i.e. tree planting) treatments, and these treatments were not applied in a consistent or systematic manner (Swan and Brown 2017: Table 2). Swan and Brown (2017) did not set a restoration criterion for site inclusion in their study, instead including all sites regardless of the combination of applied restoration treatments. I suggest that this oversight leads to unnecessary assumptions about the efficacy of restoration by assuming the effects of all treatment combinations are equivalent, and this issue could have been partially resolved a priori by hypothesizing how each treatment would affect headwater and mainstem streams and then setting requirements for site inclusion in the analyses.

**Table 1:**
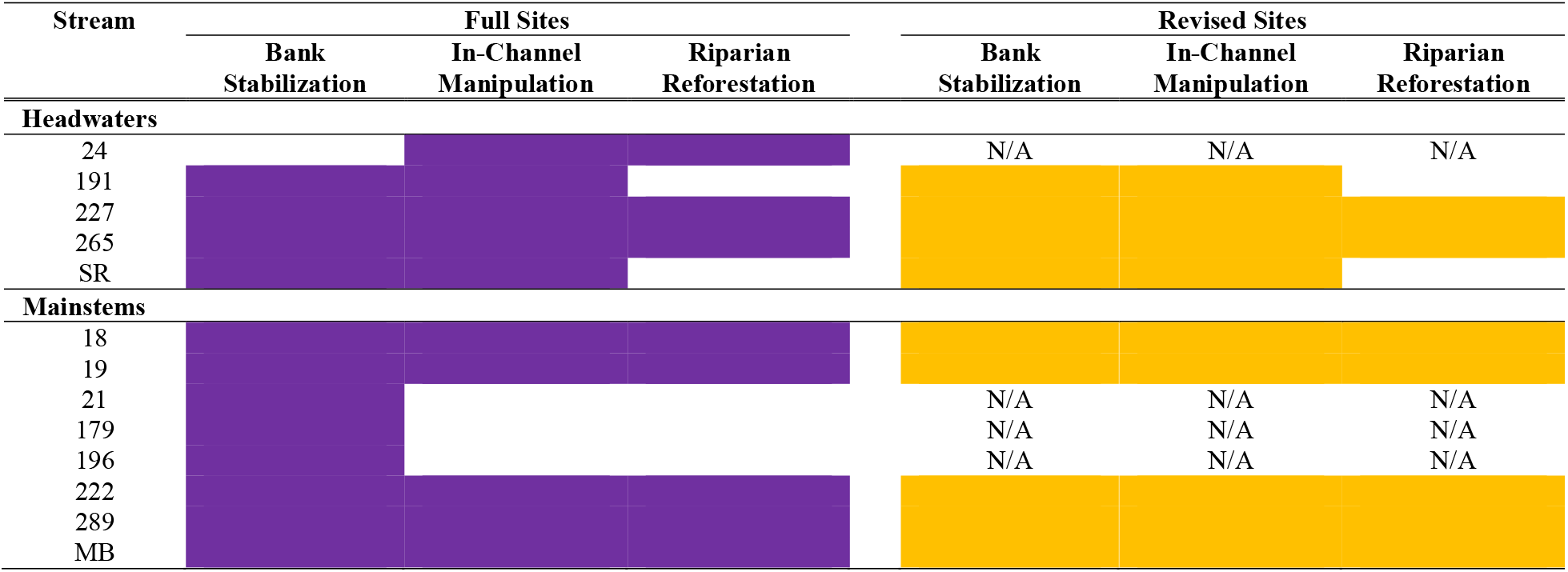
Schematic of restoration treatments applied to each stream. Shaded cells represent the restoration treatments that were applied to each stream, with full sites in purple and revised sites in yellow. Sites excluded in the revised sites analysis are denoted with N/A.

I contend that the various restoration treatments differ not only in their overall effects but also if the treatment is applied in headwater or mainstem streams, and, for these reasons, criteria for site selection could be set. I suggest that bank stabilization and in-channel manipulation treatments are more likely to have stronger and consistent effects in both headwaters and mainstems (Muotka and Syrjänen 2007, Miller et al. 2010), while riparian reforestation would likely have stronger effects in headwater compared to mainstem streams (Vannote et al. 1980, Rosi-Marshall and Wallace 2002). A similar argument was made by Swan and Brown, though it was not explicitly noted until the erratum (Swan and Brown 2018). Bank stabilization and in-channel manipulation can increase bed stability and substrate availability and diversity in both headwater and mainstem streams (Muotka and Syrjänen 2007, Miller et al. 2010), however, the effects of riparian reforestation could act on a gradient from headwaters to mainstems. For example, leaf litter is an important source of habitat and nutrients in headwaters but less so in mainstems (Vannote et al. 1980, Rosi-Marshall and Wallace 2002). Additionally, the utility of riparian reforestation on reducing nutrient inputs notwithstanding (Collins et al. 2013), the effects of riparian reforestation could be stronger in headwater streams because they are isolated systems, whereas mainstem streams receive flows of water, nutrients, and organisms from many tributaries (Vannote et al. 1980). Effectively, mainstems are dependent on other tributaries and any local restoration effects via riparian reforestation could be overwhelmed by incoming flows from unrestored streams (Wahl et al. 2013).

Here, I present a re-analysis of the data provided by Swan and Brown (2017, 2018). I hypothesized that stream-channel manipulations would have a more consistent effect between headwaters and mainstems relative to the effect of riparian reforestation, with stronger effects of restoration in headwaters relative to mainstems; I used this hypothesis to guide and inform site selection in my re-analysis. I required sites in the re-analysis to have received both the bank stabilization and in-channel manipulations treatments (hereafter “revised” sites), although sites receiving riparian reforestation were also included if they received both the bank stabilization and in-channel manipulations treatments. I also re-analyzed the full data (hereafter “full” sites) to determine if any differences, or lack thereof, between the full and revised sites analyses could be attributed to increased variation in the revised sites due to decreased sample size. Finally, I compare the interpretation and conclusions from my re-analysis to those in Swan and Brown (2017) and the erratum (Swan and Brown 2018).

## Methods

### Sampling Design

Swan and Brown (2017) conducted their study in 5 headwater and 8 mainstem streams in Baltimore County, Maryland, U.S.A. Each stream had a paired structure, where restored and adjacent, unrestored reaches were sampled; restored and adjacent reaches were separated by < 10 m. The sampling design was explicitly constructed to permit comparisons between paired restored-adjacent reaches in each of the focal streams. Each of the 13 focal streams was sampled quarterly in 2011 (spring, summer, and fall) and 2012 (winter; Swan and Brown 2017). The full sites in the re-analysis included these 13 focal streams, while the revised sites included 9 streams (4 headwaters and 5 mainstems). In the revised sites subset, 7 of the 9 sites received all three restoration treatments (i.e. bank stabilization, in-channel manipulation, and riparian reforestation); 2 of the 4 headwaters and all 5 mainstems received all restoration treatments. In contrast, 7 of the 13 streams in the full sites received all three restoration treatments: 2 of the 4 headwaters and 5 of the 8 mainstems received all restoration treatments (Table 1).

### Statistical Analyses

I generally followed the analyses as written by Swan and Brown (2017), with modifications made when necessary. The three community response variables were local diversity, spatial dissimilarity, and temporal variation. Local diversity was calculated as taxonomic richness (i.e. number of different taxa present) and taxonomic diversity (i.e. Shannon’s diversity) and compared using an analysis of variance (ANOVA). The model was constructed to examine the individual effects of reach (restored or adjacent), order (headwater or mainstem), and season (spring, summer, fall, and winter) and all two- and three-way interactions, with individual ANOVAs for richness and diversity; I also fit the full and reduced taxonomic richness models proposed in the erratum (Swan and Brown 2018) as a separate set of ANOVAs. Spatial dissimilarity between communities in restored and adjacent reaches for each order-by-season combination was quantified using the modified Gower index (Anderson et al. 2006) with a logarithm with a base of 5 on an untransformed abundance matrix. Values of the modified Gower dissimilarities were then compared using an ANOVA with the individual effects of season and order as well as their interaction. Temporal variability was measured as the multivariate dispersion (i.e. mean distance to the centroid) of repeated samples for each stream-by-reach-by-order combination (Anderson et al. 2006). Distances were calculated in principal coordinates space after Bray-Curtis dissimilarity was performed on the untransformed abundance matrix. Temporal variability values were then compared using an ANOVA with the individual effects of order and reach and their interaction. All ANOVAs were performed for both the full and revised sites, with stream identity fitted as a random effect in each ANOVA; all ANOVAs were fitted by restricted maximum likelihood.

Exploratory data analysis was conducted prior to any model fitting to determine if the data met test assumptions (Zuur et al. 2010). For the full sites analyses, numerical summaries demonstrated an unbalanced design, with equal representation of restored and unrestored reaches but a large disparity in the number of samples between headwaters and mainstems for each of the taxonomic richness and diversity (headwater n = 38, mainstem n = 62), spatial dissimilarity (headwater n = 19, mainstem = 31), and temporal variation (headwater n = 10, mainstem n = 16) analyses. Additionally, the assumption of homogeneity of variance was violated for the taxonomic richness and diversity and the spatial dissimilarity analyses. The unbalanced design was greatly reduced for the revised sites analyses: taxonomic richness and diversity (headwater n = 30, mainstem n = 40), spatial dissimilarity (headwater n = 15, mainstem = 20), and temporal variation (headwater n = 8, mainstem n = 10); however, the assumption of homogeneity of variance was still violated. To better meet the assumption of equal variance, taxonomic richness was *ln*-transformed, taxonomic diversity was square root-transformed, and spatial dissimilarity was *ln*-transformed for all analyses. Along with using transformations to response variables to better meet model assumptions, I used Type III sums of squares for evaluating main and interactive effects of factors included in the ANOVA. Swan and Brown (2017, 2018) used Type I sums of squares, which are inadequate for unbalanced and multi-factor designs with interactions between or among factors (Shaw and Mitchell-Olds 1993, Quinn and Keough 2002). Type III sums of squares are more appropriate than Type I sums of squares because: (1) tests of main effects are unweighted and unaffected by sample size; and (2) main effects are calculated after accounting for other main effects and interactions in the model, particularly when interactions are presented or hypothesized (Quinn and Keough 2002).

Model assumptions were inspected graphically, and significance was considered at *P* < 0.050. I removed the spring sample from the restored reach of site 227 from analyses because there was no corresponding sample from the adjacent reach, which would have precluded paired comparisons of restored-adjacent sites; however, I did not remove any sites prior to fitting the full and reduced model ANOVAs set by Swan and Brown (2018). Additionally, the only difference between re-analysis of the full and reduced ANOVAs set by Swan and Brown (2018) was the use of Type III sums of squares instead of Type I sums of squares. Untransformed values of variables are presented in the results and figures. All analyses were conducted using R version 3.5.3 (R Core Team 2019) with the nlme (version 3.1-139, Pinheiro et al. 2019) and vegan (version 2.5-4, Oksanen et al. 2019) packages; data and R code are deposited in the figshare repository (<u>10.6084/m9.figshare.6448010</u>). Given I made necessary modifications to the analyses written by Swan and Brown (2017, 2018), later comparisons between the re-analysis presented here and the results presented by Swan and Brown (2017, 2018) will only be in terms of statistical and ecological interpretation and not exact values of test statistics. Additionally, as the bank stabilization and in-channel manipulations were the treatments of interest for the re-analysis, later discussion of restoration will be restricted to these treatments (hereafter “channel manipulations”) unless riparian reforestation is explicitly stated. Due to the restrictions of the study design and re-analysis, riparian reforestation was a confounding treatment as bank stabilization and in-channel manipulations were the focal treatments.

To facilitate discussion among the initial study (Swan and Brown 2017), erratum (Swan and Brown 2018), and this re-analysis, effect sizes were calculated for each factor and interaction in the ANOVA models. Local diversity and temporal variability effect sizes were calculated for the erratum (Swan and Brown 2018) and full and reduced models in this re-analysis. Spatial dissimilarity effect sizes were calculated for the initial study (Swan and Brown 2017) and the full and reduced models in this re-analysis; effect sizes were not calculated for local diversity and temporal variability of the initial study (Swan and Brown 2017) as results were later corrected in the erratum (Swan and Brown 2018), and it would be illogical to make comparisons to deprecated analyses. All effect sizes were calculated as partial η^2^ (Cohen 1973):

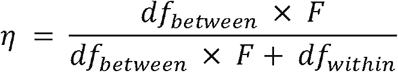

where df_between_ is the degrees of freedom associated with the factor or interaction, F is the F statistic associated with the factor or interaction, and df_within_ is the degrees of freedom associated with the residual error. Effect sizes were classified as small = 0.01, medium = 0.06, and large = 0.14 (Cohen 1973).

## Results & Discussion

There were no main or interactive effects of season, order, or reach on taxonomic richness for either the full or revised sites analyses (Table 2, Figure 1). The full model of taxonomic richness proposed in the erratum (Swan and Brown 2018) did not show any main or interactive effects of season, order, or reach (Table 3); however, the reduced model of taxonomic richness demonstrated an interaction between order and season (F_3, 80_ = 4.105, *P* = 0.009) and significant main effects of season (F_3, 80_ = 4.358, *P* = 0.007) and reach (F_1, 80_ = 4.844, *P* = 0.031). In contrast to taxonomic richness, taxonomic diversity varied by season for the full (F_3, 80_ = 12.267, *P* < 0.001) and revised (F_3, 80_ = 10.999, *P* < 0.001) sites (Table 2, Figure 2). There were no further main or interactive effects of season, order, or reach on taxonomic diversity for either the full or revised sites (Table 2, Figure 2). Spatial dissimilarity did not vary by any of the main or interactive effects of season and order for both the full and revised sites (Table 1, Figure 3). Additionally, temporal variation did not vary by the main effects of or interaction between reach and order for the full and revised sites (Table 2, Figure 4).

**Table 2:** ANOVA results for taxonomic richness, taxonomic diversity, spatial dissimilarity, and temporal variability. Models were fitted using restricted maximum likelihood and with Type III sums of squares for estimating main and interactive effects of factors.

| Source of Variation | Full Sites |  |  |  | Revised Sites |  |  |  |
| --- | --- | --- | --- | --- | --- | --- | --- | --- |
|  | numDF | denDF | F | P | numDF | denDF | F | P |
| **Taxonomic Richness** |  |  |  |  |  |  |  |  |
| Season | 3 | 73 | 1.991 | 0.123 | 3 | 47 | 1.411 | 0.251 |
| Order | 1 | 11 | 1.419 | 0.259 | 1 | 7 | 0.004 | 0.951 |
| Reach | 1 | 73 | 0.730 | 0.396 | 1 | 47 | 0.398 | 0.531 |
| Season x Order | 3 | 73 | 1.641 | 0.187 | 3 | 47 | 0.553 | 0.649 |
| Season x Reach | 3 | 73 | 0.045 | 0.987 | 3 | 47 | 0.143 | 0.934 |
| Order x Reach | 1 | 73 | 0.308 | 0.581 | 1 | 47 | 1.375 | 0.247 |
| Season x Order x Reach | 3 | 73 | 0.083 | 0.969 | 3 | 47 | 0.149 | 0.930 |
| **Taxonomic Diversity** |  |  |  |  |  |  |  |  |
| Season | 3 | 73 | 12.267 | < 0.001 | 3 | 47 | 10.999 | < 0.001 |
| Order | 1 | 11 | 2.073 | 0.178 | 1 | 7 | 0.253 | 0.630 |
| Reach | 1 | 73 | 0.088 | 0.768 | 1 | 47 | 0.085 | 0.772 |
| Season x Order | 3 | 73 | 2.477 | 0.068 | 3 | 47 | 1.272 | 0.295 |
| Season x Reach | 3 | 73 | 0.323 | 0.809 | 3 | 47 | 0.284 | 0.837 |
| Order x Reach | 1 | 73 | 0.139 | 0.710 | 1 | 47 | 0.917 | 0.343 |
| Season x Order x Reach | 3 | 73 | 0.313 | 0.816 | 3 | 47 | 0.149 | 0.930 |
| **Spatial Dissimilarity** |  |  |  |  |  |  |  |  |
| Season | 3 | 31 | 1.146 | 0.346 | 3 | 20 | 0.425 | 0.737 |
| Order | 1 | 11 | 0.062 | 0.808 | 1 | 7 | 0.007 | 0.934 |
| Season x Order | 3 | 31 | 1.356 | 0.275 | 3 | 20 | 0.306 | 0.821 |
| **Temporal Variability** |  |  |  |  |  |  |  |  |
| Order | 1 | 11 | 0.030 | 0.866 | 1 | 7 | 0.957 | 0.361 |
| Reach | 1 | 11 | 1.906 | 0.195 | 1 | 7 | 2.693 | 0.145 |
| Order x Reach | 1 | 11 | 2.054 | 0.180 | 1 | 7 | 4.718 | 0.066 |

**Figure 1:**
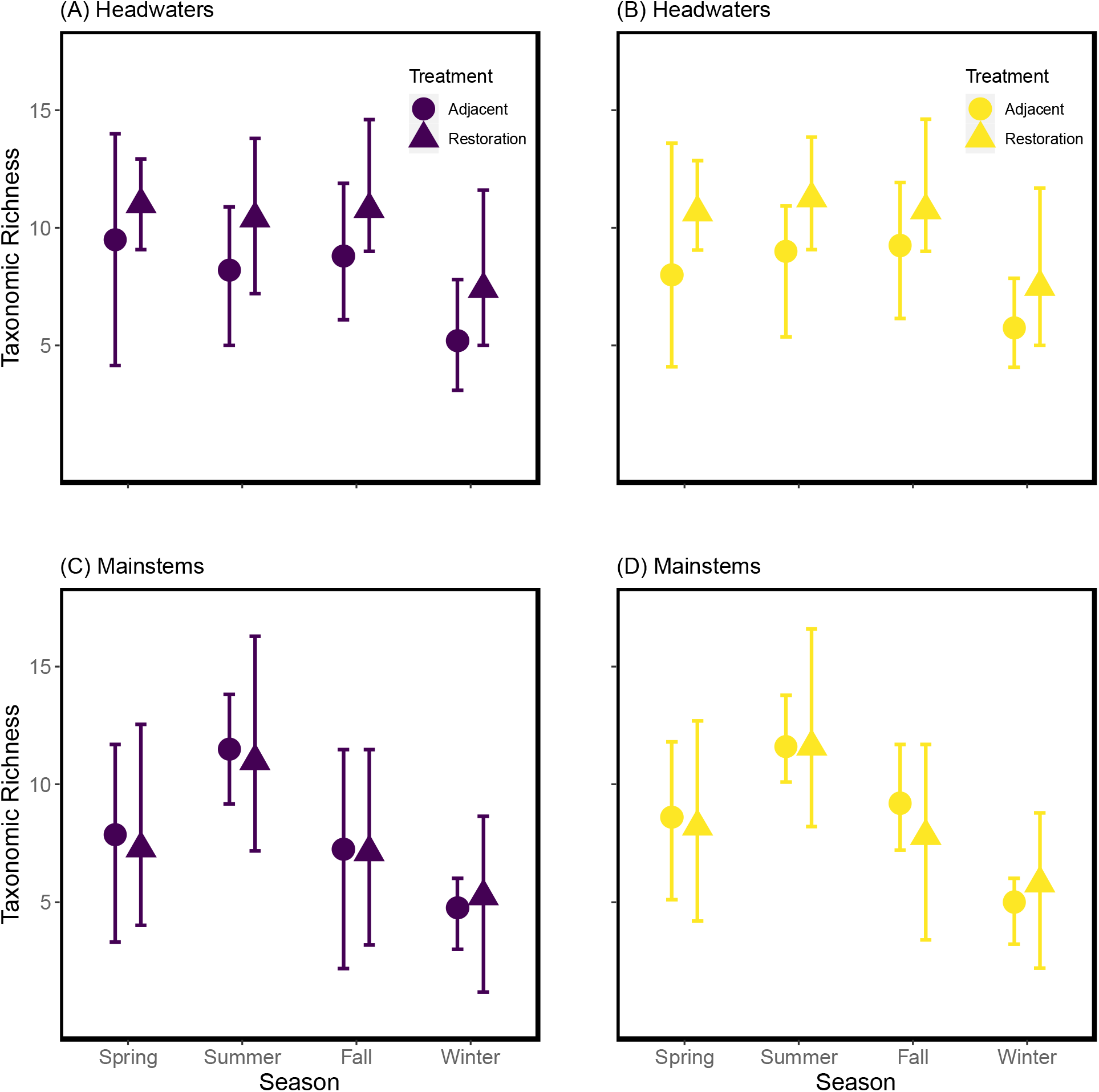
Plots of taxonomic richness in headwaters (A and B) and mainstems (C and D). Taxonomic richness is reported for restored (triangles) and adjacent (circles) sites; values from the full sites are reported in purple (A and C), while those from the revised sites are reported in yellow (B and D). Points represent mean ± 95% CI.

**Table 3:** ANOVA results for the full and reduced models of taxonomic richness proposed by Swan and Brown (2017, 2018). Models were fitted using restricted maximum likelihood and with Type III sums of squares for estimating main and interactive effects of factors. The unpaired sample was not removed from the data prior to fitting the full and reduced model ANOVAs.

| Source of Variation | Full Model |  |  |  | Reduced Model |  |  |  |
| --- | --- | --- | --- | --- | --- | --- | --- | --- |
|  | numDF | denDF | F | P | numDF | denDF | F | P |
| Season | 3 | 74 | 2.478 | 0.068 | 3 | 80 | 4.360 | 0.007 |
| Order | 1 | 11 | 0.925 | 0.357 | 1 | 11 | 1.579 | 0.235 |
| Reach | 1 | 74 | 1.409 | 0.239 | 1 | 80 | 4.844 | 0.031 |
| Season x Order | 3 | 74 | 2.359 | 0.078 | 3 | 80 | 4.105 | 0.009 |
| Season x Reach | 3 | 74 | 0.152 | 0.928 | N/A | N/A | N/A | N/A |
| Order x Reach | 1 | 74 | 0.979 | 0.326 | 1 | 80 | 3.530 | 0.064 |
| Season x Order x Reach | 3 | 74 | 0.071 | 0.976 | N/A | N/A | N/A | N/A |

Note: N/A indicates a factor or interaction that was removed in the reduced model.

**Figure 2:**
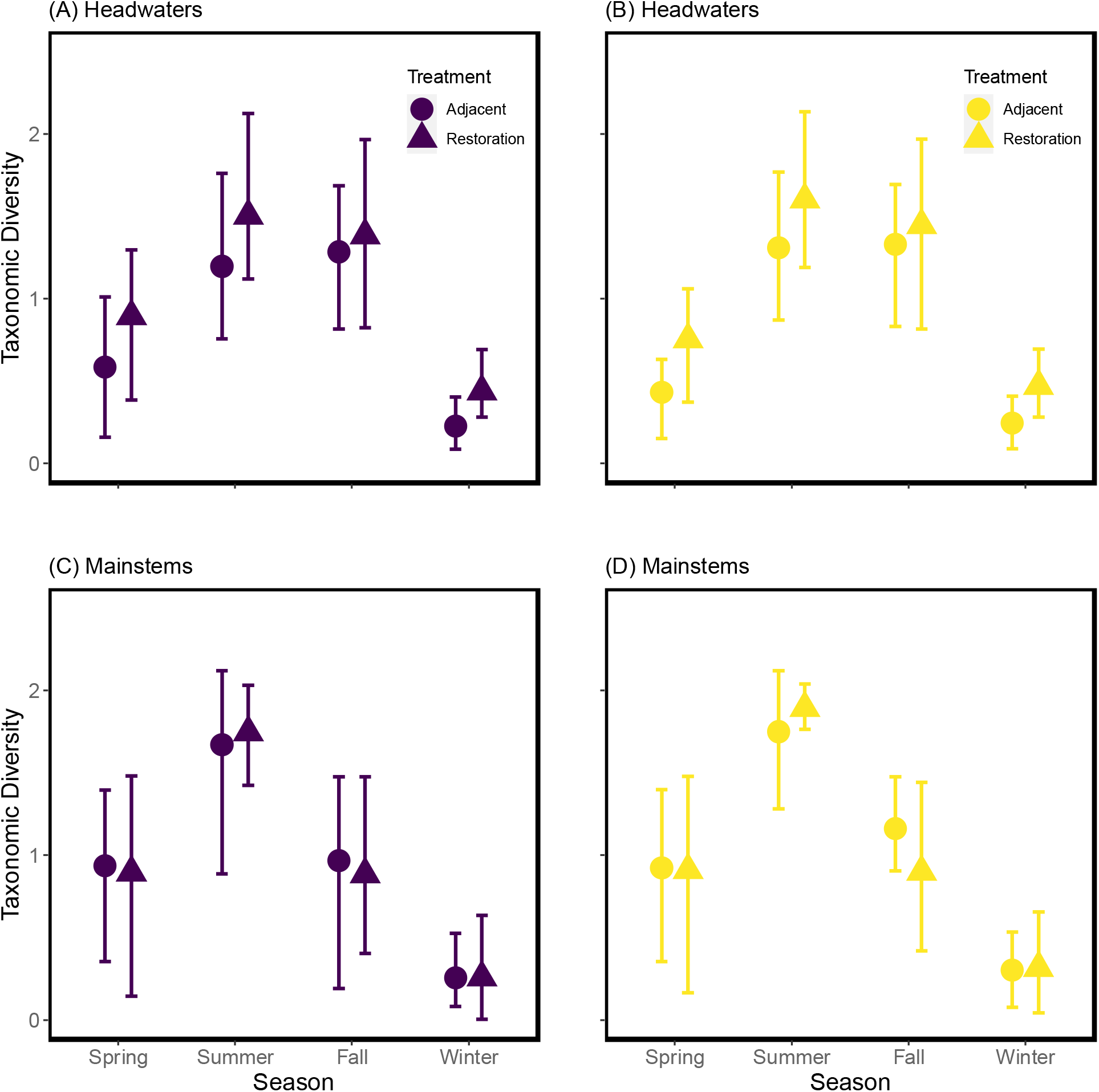
Plots of taxonomic diversity, calculated as Shannon’s diversity, in headwaters (A and B) and mainstems (C and D). Taxonomic diversity is reported for restored (triangles) and adjacent (circles) sites; values from the full sites are reported in purple (A and C), while those from the revised sites are reported in yellow (B and D). Points represent mean ± 95% CI.

**Figure 3:**
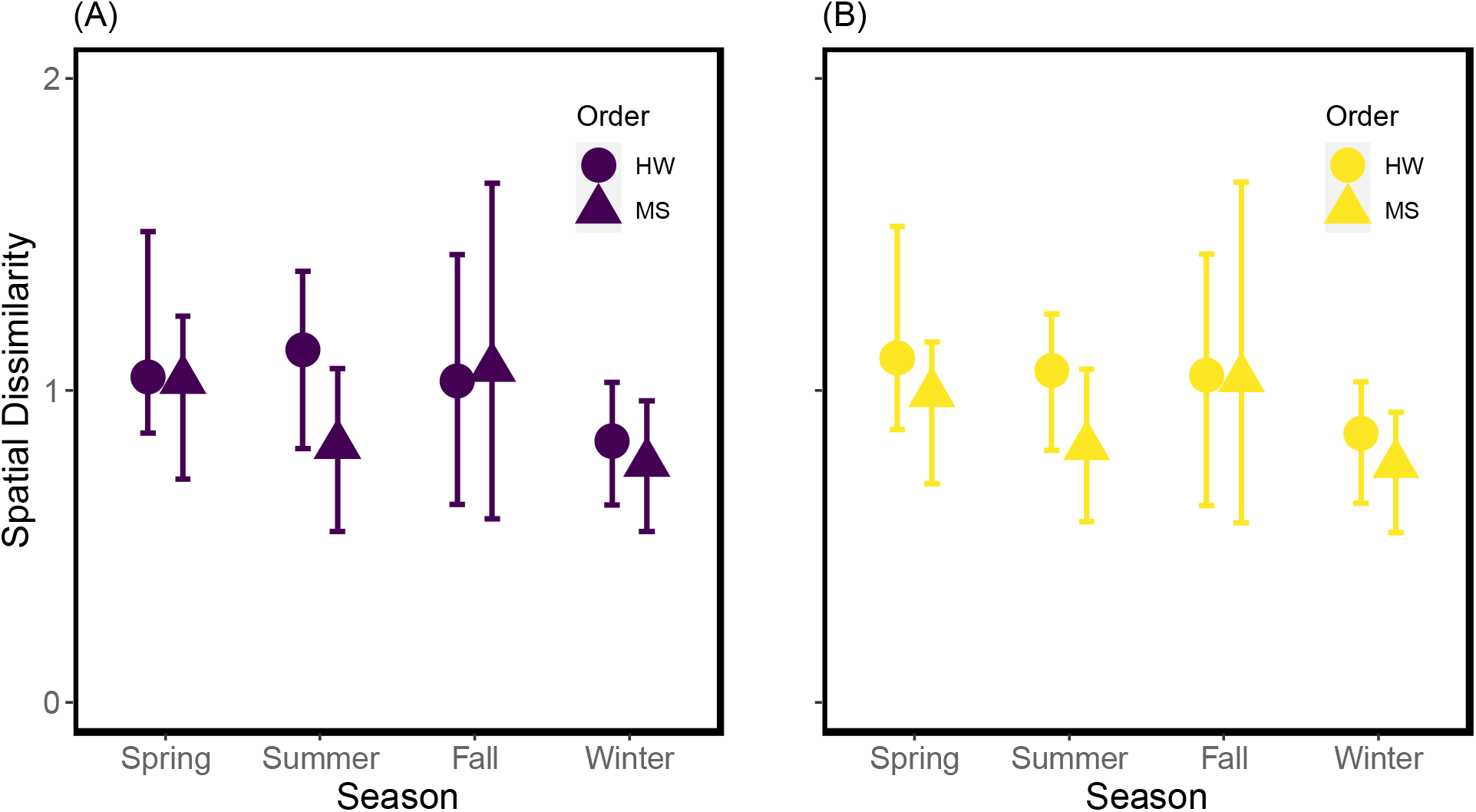
Plots of spatial dissimilarity between paired restored and adjacent reaches. Estimates are reported for headwaters (HW, circles) and mainstems (MS, triangles); values from the full sites are reported in purple (A), while those from the revised sites are reported in yellow (B). Points represent mean ± 95% CI.

**Figure 4:**
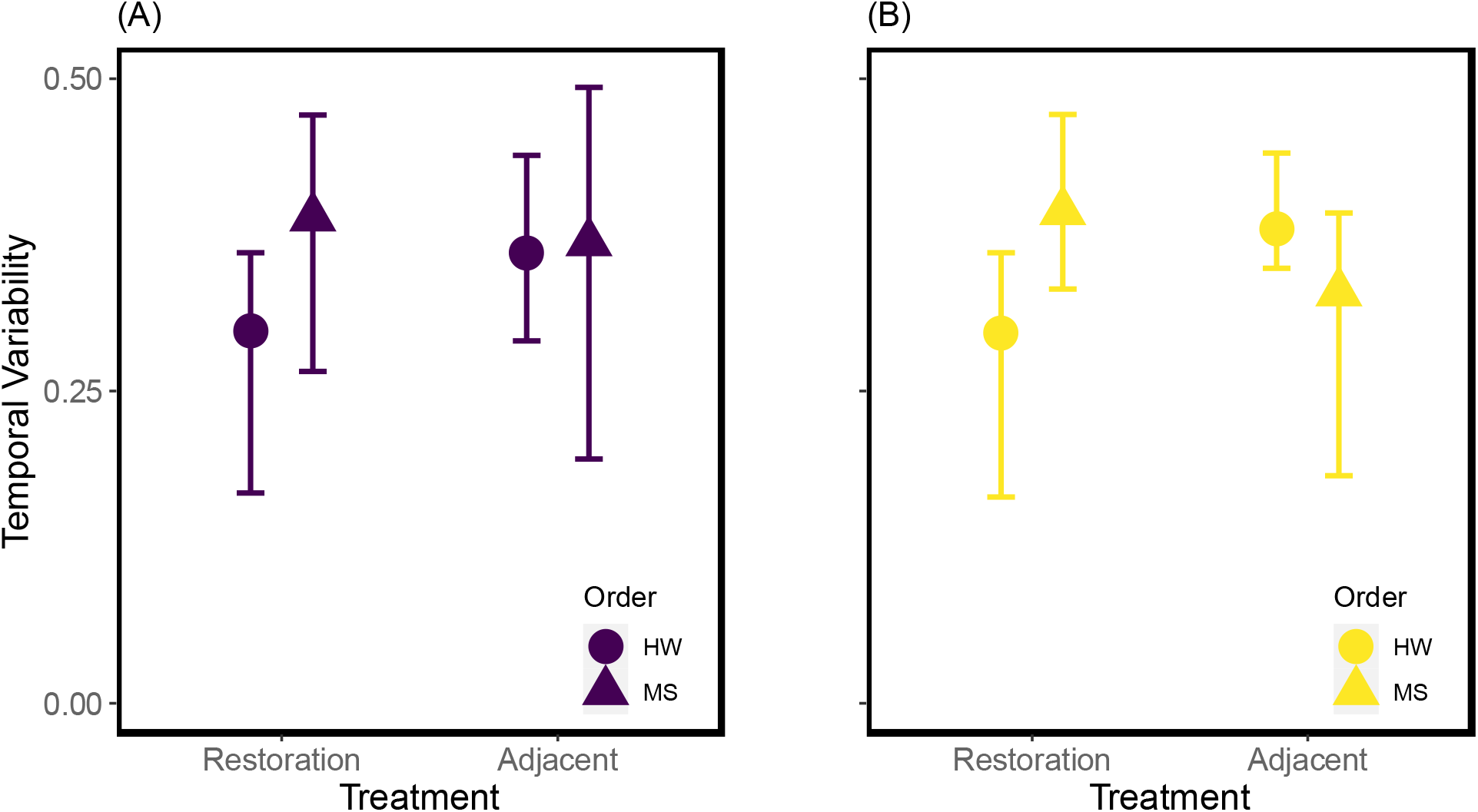
Plots of temporal variability. Estimates are reported for headwaters (HW, circles) and mainstems (MS, triangles); values from the full sites are reported in purple (A), while those from the revised sites are reported in yellow (B). Points represent mean ± 95% CI.

Analyses by Swan and Brown (2017, 2018) overestimated some effect sizes for the local diversity and spatial dissimilarity analyses (Table 4). Large effect sizes were observed for season in the local diversity (η^2^ = 0.1405) and spatial dissimilarity analyses (η^2^ = 0.1817) by Swan and Brown (2017, 2018), but there was a medium effect size for season for the full (η^2^ = 0.0756) and revised (η^2^ = 0.0826) sites in the re-analysis of local diversity. A medium effect size for season was observed for the full sites (η^2^ = 0.0999) and a small effect size was observed for the revised sites (η^2^ = 0.0599) for the spatial dissimilarity analyses. Similar effect sizes for order for the local diversity analyses were observed for the Swan and Brown (2017, 2018) analyses (η^2^ = 0.1255) and the full sites (η^2^ = 0.1143) analysis, but the revised sites analysis had a negligible effect size for order (η^2^ = 0.0006). Noticeably, equivalent and small effect sizes of reach were observed for the local diversity analyses across in the re-analysis but not in the original study (Swan and Brown 2017, 2018; Table 4). In contrast to local diversity and spatial dissimilarity, Swan and Brown (2017, 2018) frequently underestimated effect sizes for the temporal variability analyses (Table 3). A medium effect size for order (η^2^ = 0.1203) and large effect sizes for reach and the order-by-reach interaction were observed in the revised sites analysis (reach η^2^ = 0.2779, order-by-reach η^2^ = 0.4026), with the largest effect size from all analyses and community diversity metrics derived from the order-by-reach interaction in the revised sites analysis (η^2^ = 0.4026). Although this effect was not statistically significant *(p* = 0.066), it suggests that channel manipulation treatments could have an effect that is dependent on network position, but the statistical power was insufficient.

**Table 4:**
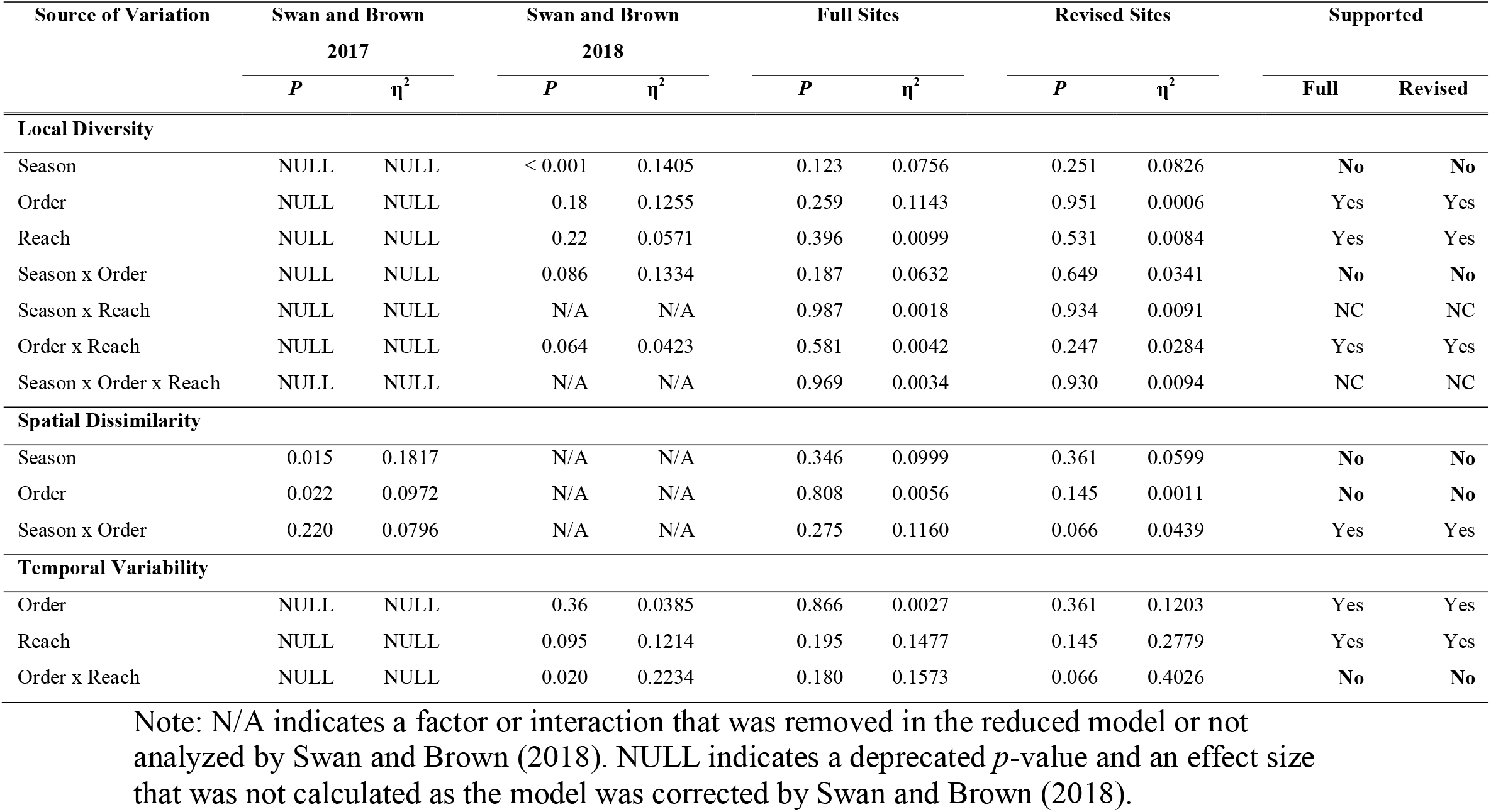
Comparison of ANOVA results and effect sizes (η^2^) between the initial study (Swan and Brown 2017), the erratum (Swan and Brown 2018), and if results are consistent upon re-analysis in the full (Full) or revised (Revised) sites analyses. Support is denoted as: Yes = consistent with both the initial study and the erratum; No = inconsistent with both the initial study and erratum; NC = no comparisons can be made as the factor or interaction is missing from the previous analysis. Spatial dissimilarity was not re-analyzed in the erratum, so results are only provided for the initial study. Bold values in the Supported column indicate differences in statistical significance between the initial study and/or erratum and the re-analysis.

Differences in significant main effects or interactions within the full and revised sites in the re-analysis did not seem to be the result of increased variation in the revised sites. In fact, variance, as measured by 95% confidence intervals, was either similar or even reduced for each of local diversity, spatial dissimilarity, and temporal variability for the revised sites compared to the full sites (Figures 1–4). In general, local diversity more frequently increased in headwaters relative to mainstems (Figure 5), and this trend was consistent for both the full (Figure 5A and 5B) and revised (Figure 5C and 5D) sites. Taken together, it is unlikely that that revised sites analysis was unable to detect effects due to increased variation and more likely due to reduced statistical power associated with a smaller sample size or the true lack of an effect of channel manipulations on different facets of biodiversity in this system.

**Figure 5:**
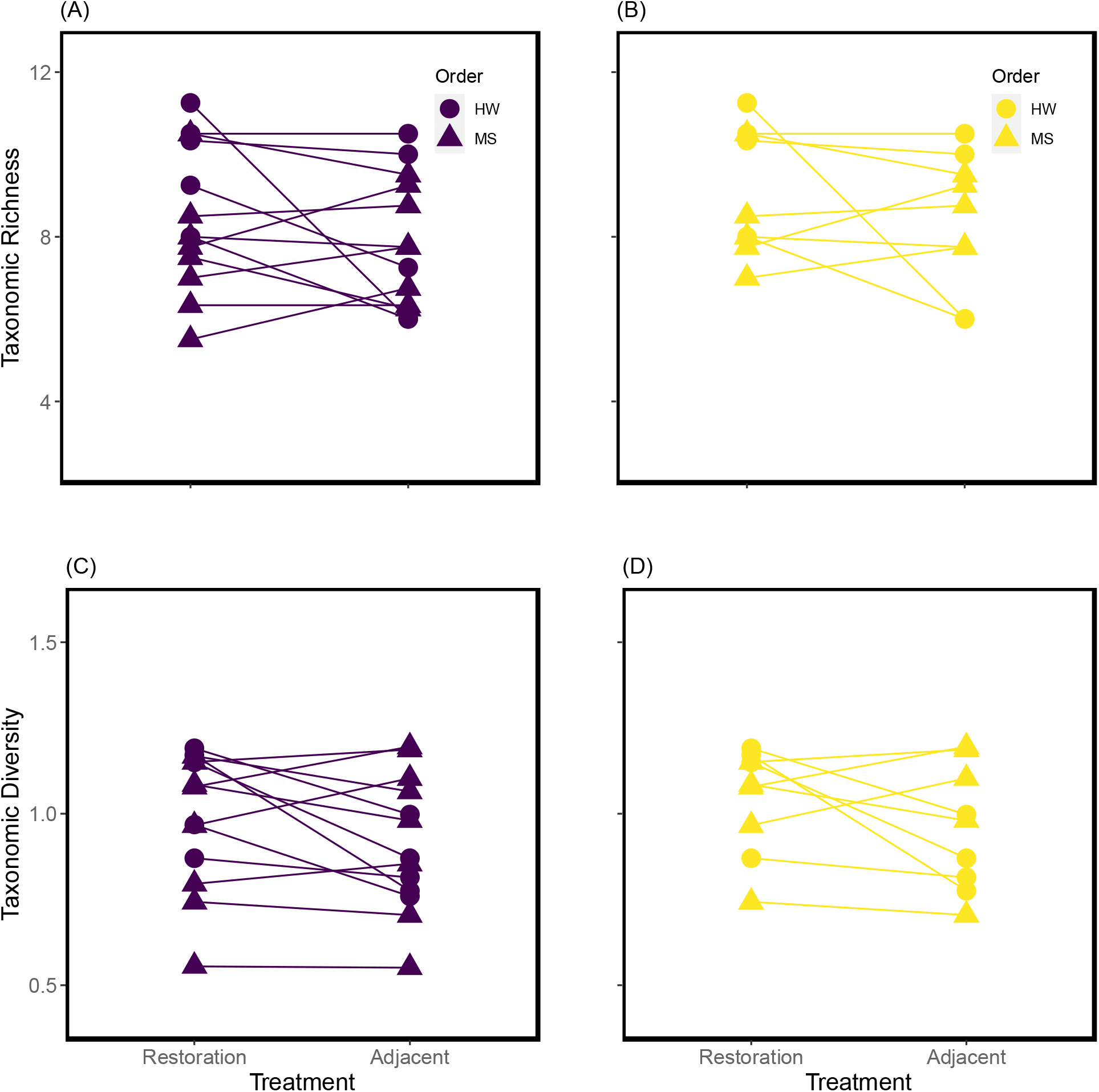
Plots of taxonomic richness (A and B) and diversity (C and D) for restored and adjacent reaches of each site. Lines connect values for each site. Values from the full sites are reported in purple (A and C), while values from the revised sites are reported in yellow (B and D). Points represent the mean taxonomic richness or diversity. Error bars are not presented to improve graphical clarity.

### Effectiveness of Local Restoration

I hypothesized that channel manipulations would have stronger effects in headwaters relative to mainstems. As there were no significant effects of channel manipulations on any of the community metrics between headwaters and mainstems, this hypothesis could be invalid or, at a minimum, revised and re-tested in channel manipulations experiments. I was unable to directly test this hypothesis because I was re-analyzing data from a previous study and the experimental design precluded any test to isolate the effects; however, the hypothesis was intended to guide criteria for site selection and reduce variation in restoration treatments among sites and not to necessarily or strictly compare the effects channel manipulations and riparian reforestation treatments on biodiversity in restored streams. Despite these limitations, there is some evidence of a large effect in the revised sites on temporal variation in community composition, with this effect dependent on network position. The re-analysis was lacking sufficient statistical power to combine statistical significance with ecological relevance, potentially due to inconsistencies in applied restoration treatments. Only the mainstem stream group received all channel manipulation and riparian reforestation treatments, while the riparian reforestation treatment was applied to half headwater stream group (2/4 streams; Table 1). As headwater streams are strongly linked to allochthonous inputs and leaf litter subsidies from the riparian forest for community structure (Wallace et al. 1997), the lack of riparian reforestation to all streams in the headwater group could have reduced any observable recovery of biodiversity. More consistent application of all restoration treatments across both headwater and mainstem streams could provide the conditions for more effective restoration and recovery of stream ecosystems, measured by local diversity, spatial dissimilarity, and temporal variation. Based on the evidence from this re-analysis, further evaluation of this interactive effect of restoration and network position on temporal variability could be an efficacious avenue for bridging metacommunity ecology and restoration efforts in rivers and streams and increasing positive biodiversity outcomes.

Regarding evidence for stream-channel manipulations and other treatments for effective restoration, previous research suggests local habitat manipulations are ineffective for structuring communities and increasing biodiversity (Palmer et al. 2010). An emerging hypothesis is that local factors, such as habitat complexity and water quality, are overwhelmed by regional factors, such as dispersal and position within the larger network (Heino 2013, Tonkin et al. 2014). Given channel manipulations did not have a statistically-significant effect on any diversity measure of communities in either headwaters or mainstems and that the majority of effect sizes were small-to-medium (Table 3), this could suggest either channel manipulations were either wholly inadequate for both headwaters and mainstems or that the larger network and regional species pool were already degraded (Sundermann et al. 2011), overwhelming any mitigating effects of channel manipulations.

### Restoration Ecology & Experimental Design

Restoration of the streams was done in isolation of the study design and prior to data collection, resulting in variation in the types of treatments applied to the streams (Swan and Brown 2017). Although Swan and Brown (2017) noted this limitation of their study, they did not acknowledge they could have better controlled for this variation by setting strict criteria for site selection and inclusion, which informed my hypothesis and was the foundation for my re-analysis. This concern was briefly acknowledged in the erratum (Swan and Brown 2018), where the data quality control process removed sites if they only received riparian reforestation treatments without at least one of either the bank stabilization or in-channel manipulation treatments; however, Swan and Brown (2017, 2018) proceeded to analyze data from sites receiving any resulting combination of restoration treatments, despite suggesting that in-stream modification treatments would have stronger effects on communities relative to riparian reforestation (Swan and Brown 2018). Setting a more stringent criterion for site inclusion, as was done in this re-analysis of the revised sites, would have reduced the variation in the applied restoration treatments and provided a more balanced experimental design.

The inconsistent application of restoration treatments prohibited a robust evaluation that could have been possible with a factorial experiment; therefore, the singular and interactive effects of the restoration treatments in the study system remain untested. This further complicates the indiscriminate usage of “restoration” by Swan and Brown (2017, 2018) as the underlying mechanism of restoration on the stream invertebrate communities remains an unknown quantity. Identifying how individual and combinations of restoration treatments affect stream communities would provide valuable insight for maximizing the effectiveness of restoration efforts. In the absence of this knowledge, reducing the variation in which restoration treatments were applied to the streams, as done with revised sites analysis, arguably would have been a better avenue. Additionally, restoration treatments were not applied to all sites at the same time, which could allow for further confounding variation among sites. Previous research has demonstrated mixed results of the effects of time since restoration on biodiversity responses (Miller et al. 2010, Orzetti et al. 2010, Louhi et al. 2011), but it is hypothesized that more time allows for increased colonization of restored habitats and for restoration treatments (e.g. riparian reforestation) to have an impact on the system (Lake et al. 2007, Palmer et al. 2014). Knowledge of the effects of restoration treatments, both in isolation and in combination, and incorporating time since restoration in evaluations of community responses to restoration would likely improve future studies and experiments.

### Statistical Inconsistencies

Channel manipulations were not found to have a significant effect on local diversity, spatial dissimilarity, or temporal variability of stream invertebrate communities between paired restored and unrestored reaches in headwaters and mainstems. These results presented here, not exact values of test statistics but in terms of interpretation, contradict the results presented in the original paper (Swan and Brown 2017) and in the erratum (Swan and Brown 2018; Table 3). This is concerning, as any data management and analytical errors in the original paper were supposedly resolved in the erratum (Swan and Brown 2018); however, the discrepancies can be partially explained by the erroneous reporting and implementation of statistical analyses. First, and as was noted above, Swan and Brown (2017, 2018) analyzed an unbalanced design with unequal variance using an ANOVA with Type I sums of squares, when transformations to response variables were necessary to better meet test assumptions and Type III sums of squares were more appropriate for investigating the main and interactive effects (Shaw and Mitchell-Olds 1993, Quinn and Keough 2002). Second, the fitting of the random effects in the ANOVAs was incorrect. Swan and Brown (2018) reported fitting stream identity as a random effect. With the R code provided with the erratum (Swan and Brown 2018: Supporting Information), each site-by-reach combination was fitted as a random effect, despite adjacent and restored reaches being separated by < 10 m (Swan and Brown 2017). Fitting stream identity alone would have been a more appropriate fitting of a random effect that reflected the non-independence of the reaches at such small spatial scales, and, importantly, an accurate implementation of the written methods. Additionally, with the evidence available, only the local diversity analysis had a random effect fitted with the model, despite a random effect being applicable to all analyses based on the experimental design. Third, removal of the restored site without a paired observation is necessary for the fundamental goal of the study: comparing community diversity between paired restored and adjacent reaches across headwater and mainstem streams. Without removing the site, comparisons would be made to an unpaired reach, violating the experimental design and central goal of the study.

Finally, there was disagreement between the reported analytical procedure and what was actually conducted when analyzing temporal variability. Temporal variability was reportedly quantified as the mean distance to the group centroid (Anderson et al. 2006) after applying a Bray-Curtis dissimilarity index on an untransformed abundance matrix (Swan and Brown 2017). Results presented in the erratum were actually derived from the spatial median after a Jaccard index was applied to a presence-absence matrix (Swan and Brown 2018: Supporting Information); in the initial study, Swan and Brown (2017) stated that a Jaccard index applied to a presence-absence matrix would produce stronger results than a Bray-Curtis index applied to an abundance matrix. No random effect of stream identity was fitted for this ANOVA, although it would have been appropriate given the study design (Quinn and Keough 2002). Most importantly, none of the changes to the analytical procedure were reported in the erratum, and these alterations were only found upon evaluation of the provided R code (Swan and Brown 2018: Supporting Information). Without consulting the supporting information or if no R code was provided, it would have been assumed the results presented in the erratum (Swan and Brown 2018) were derived from the analytical procedure described in the original study (Swan and Brown 2017), just with the corrected dataset; this assumption would have been incorrect.

### Ecological Implications & Prospective Suggestions

Based on my re-analysis, I have concluded that, given channel manipulations had no effect on any community diversity metric, local restoration of streams can be ineffective if (1) both dispersal and habitat quality are structuring biodiversity (Heino et al. 2015, Smith et al. 2015, Downes et al. 2017) and (2) the larger regional community is already degraded (Sundermann et al. 2011). Future research projects should experimentally- and factorially-manipulate different environmental factors to evaluate the relative impact and effectiveness of restoration techniques, which could allow for the identification of how in-stream and riparian reforestation treatments affect stream biodiversity. More broadly, restoration of individual reaches of rivers and streams might be insufficient to reach objectives of restoration projects (Bernhardt and Palmer 2011, Sundermann et al. 2011). Streams and rivers will likely have better restoration success if a regional approach is taken (Bernhardt and Palmer 2011), whereby heterogeneity among rivers and streams in ecological, geographic, and spatial context is incorporated (Palmer et al. 2010, 2014, Booth et al. 2016). In this respect, I agree with Swan and Brown (2017, 2018), along with previous research, that local manipulations are insufficient, and a regional perspective is needed for more effective restoration in rivers and streams.

## Acknowledgements

I am grateful for the comments and discussion provided by Karl Cottenie, Mariana Perez Rocha, and Eric Harvey that greatly improved the quality of the manuscript.

## Data Accessibility

Data and R code are deposited in the figshare repository (<u>10.6084/m9.figshare.6448010</u>).

